# Experimental-based Computational Modeling Distinguishes Early Cardiac Outflow Tract Compensation Mechanisms

**DOI:** 10.1101/2020.09.11.292714

**Authors:** Stephanie E. Lindsey, Irene E. Vignon-Clementel, Jonathan T. Butcher

## Abstract

Mechanical forces are essential for proper growth and remodeling of the primitive pharyngeal arch arteries (PAAs) into the great vessels of the heart. Despite general acknowledgement of a link between abnormal hemodynamics and cardiac malformations, the direct correlation between hemodynamics and pharyngeal arch artery morphogenesis remains poorly understood. The elusiveness behind understanding hemodynamic-malformation links is largely due to the difficulty of performing isolated hemodynamic perturbations and quantifying key hemodynamic indices in-vivo. To overcome this issue, minimally invasive occlusion experiments were combined with three-dimensional anatomical models of development and in-silico testing of experimental phenomenon. This combined experimental-computational approach led to a mechanistic understanding of physiological compensation mechanisms in abnormal cardiac morphogenesis. Using our experimental-based framework, we detail morphological and hemodynamic changes twenty-four hours post vessel occlusion. To gain mechanistic insights into the dynamic vessel adaptation process, we perform in-silico occlusions which allow for quantification of instantaneous changes in mechanical loading. We follow the propagation of small defects in a single embryo Hamburger Hamilton (HH) Stage 18 embryo to a more serious defect in an HH29 embryo. Results demonstrate that abnormal PAA hemodynamics can precipitate abnormal cardiac function given the correct timing and location of injury. Following vessel occlusion, morphology changes along the arches are no longer a simple flow-mediated response but rather work to maintain a range of wall shear stress values. Occlusion of the presumptive aortic arch overrides natural growth mechanisms and prevents it from becoming the dominant arch of the aorta.

**Author Summary:** The developing great vessels transport flow from the heart to the rest of the body. Proper spatial temporal morphogenesis of the primitive paired vessels into the definitive outflow tract of the heart is critical for normal cardiac function. Malpatterning of the great vessels is highly prevalent in congenital heart defects and occurs in conjunction with other intracardiac malformations, such as impaired ventricle and valve development. In this work, we combine experimental-based computational modeling with theoretical adaptation principles. Our combined experimental-computational framework allows for the delineation of immediate and longer-term vascular remodeling as well as the physical mechanisms behind such changes. We show that a small flow obstruction originating within the developing vessels can propagate into structurally serious malformations with impaired functionality.

## Introduction

During cardiac morphogenesis, blood exits the developing chick embryo’s heart through the pharyngeal arch artery system. These six bilateral paired vessels exist in various combinations as they sequentially emerge, remodel and disappear before forming the mature aortic arch, pulmonary artery, pulmonary veins and venae cavae (Figure 1). Hemodynamics plays a vital role in the maturation of the pharyngeal arch artery system [1]–[4]. The role of abnormal hemodynamics in the creation of cardiac abnormalities has intrigued researchers for decades [4]–[12]. Disruption of established flow patterns during critical windows of development produces a range of defects that drastically alter function of the mature heart. These defects may stem from improper cardiac looping, incomplete outflow tract rotation, incomplete outflow tract septation, abnormal maturation of the cardiac valves or yet to be determined factors. Malformations of the outflow tract account for over 50% of clinically relevant congenital heart defects [13]. Determining the origin of such defects continues to pose a major challenge to the field. Cardiac morphogenesis is a complex interconnected process that is difficult to delineate. Altered hemodynamic flow patterns in the heart following surgical manipulation of the atrium, have been shown to affect the ventricle, as well as the pharyngeal arch artery system. Ligation of the left atria redistributes flow to the right side of Hamburger Hamilton (HH) Stage 21 embryos, immediately affecting flow patterns and hemodynamic forces in the arch arteries. Within 24 hours of left atrial ligation, the fourth aortic arch of HH24 embryos is drastically reduced in dimensions. By HH27 (48 hours post ligation) arches were skewed and distorted. All embryos surviving to HH34 (day 8) displayed hypoplastic ventricles; 70% also possessed abnormal aortic arch patterning [4].

**Figure 1:**
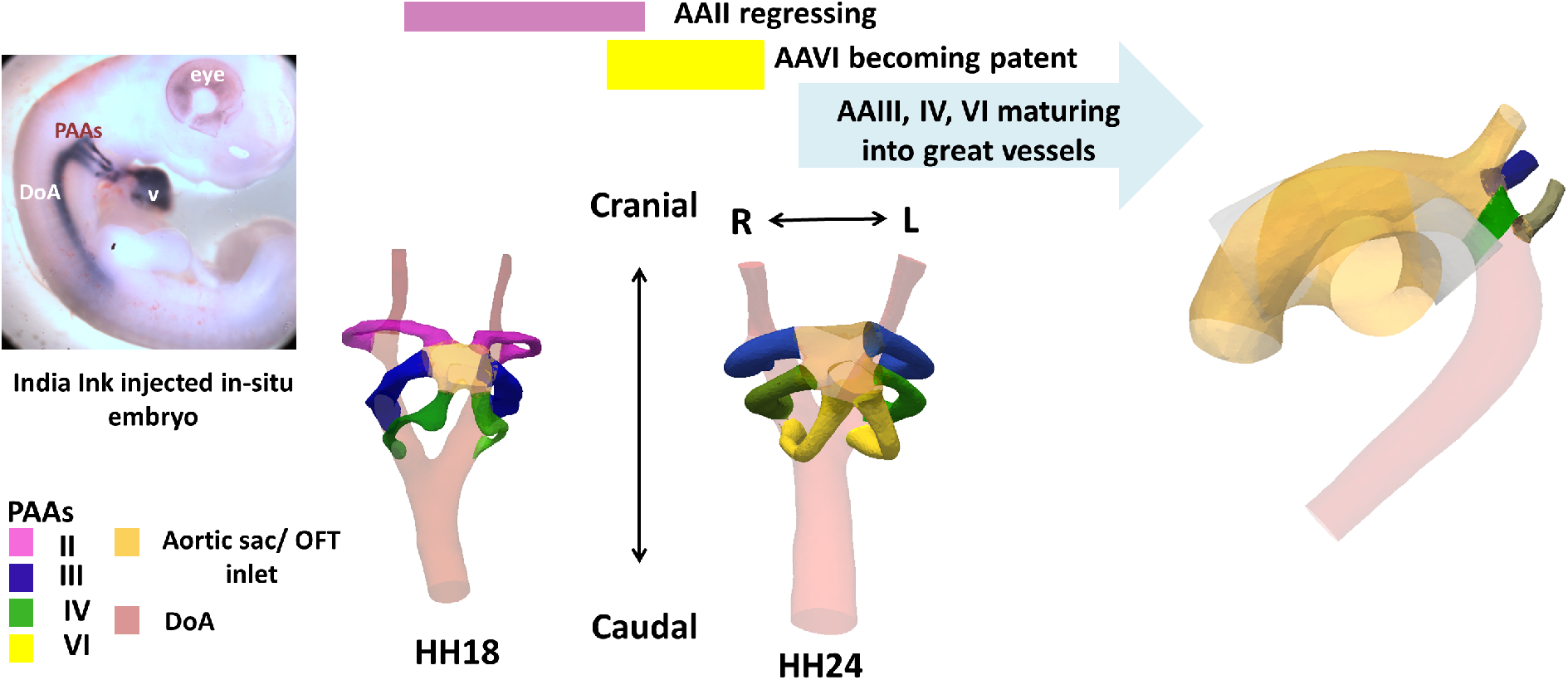
Anatomical Reference Figure. Summary of arches present per stage and their contributions to the final aortic arch configuration (arches not shown to scale).

While much has been gained from ligation experiments such as those performed by Hu et al., a major limitation in determining the causality of clinically relevant cardiac abnormalities is the difficulty of studying the effects of hemodynamics alone in the creation of cardiac abnormalities. In an effort to surgically manipulate flow patterns in experimental animal models, researchers have used a combination of ligations, clipping, and cauterizations, all of which disrupt flow patterns, as well as change the properties of the surrounding tissues. Alteration of mechanical stress and strains in the developing heart have been shown to regulate vascular growth and remodeling as well as trigger cell signaling and subsequent organization patterns [12], [14], [15]. Recent advances in imaging technology have allowed for the minimally invasive occlusion of flow in the developing vascular system [3]. Studies such as these may prove to be important in delineating the sequence of events that leads to clinically relevant cardiac abnormalities. The coupling of computational modeling with targeted experimental studies can provide further insight into the mechanisms behind abnormal cardiac morphogenesis, correlating sites of altered wall shear stress (WSS) with commonly affected areas in the clinically associated disease model [2] or highlight driving forces in development [3], [16].

Another major limitation in determining causality for cardiac malformations is an inability to create, study or identify what likely begins as a subtle abnormality before propagating into clinically serious malformations. For this reason, we characterized normal PAA development in a cohort-based study that identified a range of normal development in Hamburger Hamilton (HH) staged 18, 24, and 26 embryos, taking into account morphological and hemodynamic parameters and determining a relationship between the two [17]. In this study, we examine the effects of vessel occlusion on PAA development as a biological model for outflow tract malformations resulting from abnormal PAA development. As full vessel occlusion has been shown to be particularly lethal [3], here we focus on partial occlusions which are likely more representative of clinically serious congenital heart defects where the fetus is carried to term but the heart is malformed. The word ‘occlusion’ is used to refer to both full and partial vessel occlusions, with the distinction being made when necessary. Using detailed 3D reconstructions of experimental occlusion geometries, we analyze vessel adaption to abnormal hemodynamic conditions that originate with the arch arteries themselves. We compare experimental results post-occlusion to that of in-silico occlusions taken at the time of intervention (HH18) and 24-hours post intervention (HH24) when the embryo’s vasculature has had time to remodel and adapt. We subsequently demonstrate how an embryo’s “small deviation from normal” can propagate into a more serious malformation. Ultimately this work details structural compensation mechanisms involved in early cardiac outflow morphogenesis and the effects on hemodynamic function and subsequent cardiac growth.

## Results

### Early great vessel morphogenesis depends on hemodynamic flow patterns

In order to explore embryo compensation mechanisms following altered hemodynamic flow, we occluded the developing IVR PAA in HH18 embryos using nonlinear optical techniques that allowed for changes in hemodynamics alone (see methods). To date, this is the only experimental method that locally and noninvasively alters flow patterns. Occlusion experiments were purposely performed on HH18 embryos and selected to have a narrow (nascent) PAA IV vessel in order to facilitate vessel occlusion. Although PAA IVR was small (~30 μm in diameter) at time of occlusion(HH18), by HH24 it carries approximately 22% of flow exiting the heart [17].

Following HH18 vessel occlusion, the largest percentage of flow was re-directed to PAA IIIL in the partial occlusion geometries and PAA VIL in the case of full vessel occlusion, with PAA IIIL carrying 35% of flow compared to 20% of flow in its control counterpart in partial occlusion geometries and PAA VIL carrying 31% of flow compared to 7% in its control counterpart in full occlusion geometries (Figure 2A). Within 24 hours post-occlusion, embryos exhibited a range of structural defects (Supplemental Figures 1,2). Observed malformations include the merging of arch arteries into a single vessel before separating into two distinct vessels, abnormal arch artery spacing, skewed right and left branches, enlarged arch arteries and outflow inlets, abnormal rotation of the outflow tract (OFT) junction, abnormal bulging of the aortic sac and abnormal patterning of the cranial DoA branches. Position and length of the initial vessel occlusion were associated with certain types of defects. Occlusions occurring near the DoA junction were associated with merged vessel pathways originating from the aortic sac, while occlusions occurring near the aortic sac junction were associated with enlarged OFT orifices. More elongated vessel occlusions, that spanned more of the lateral length, were associated with abnormal spacing and skewed left and right patterning.

**Figure 2.**
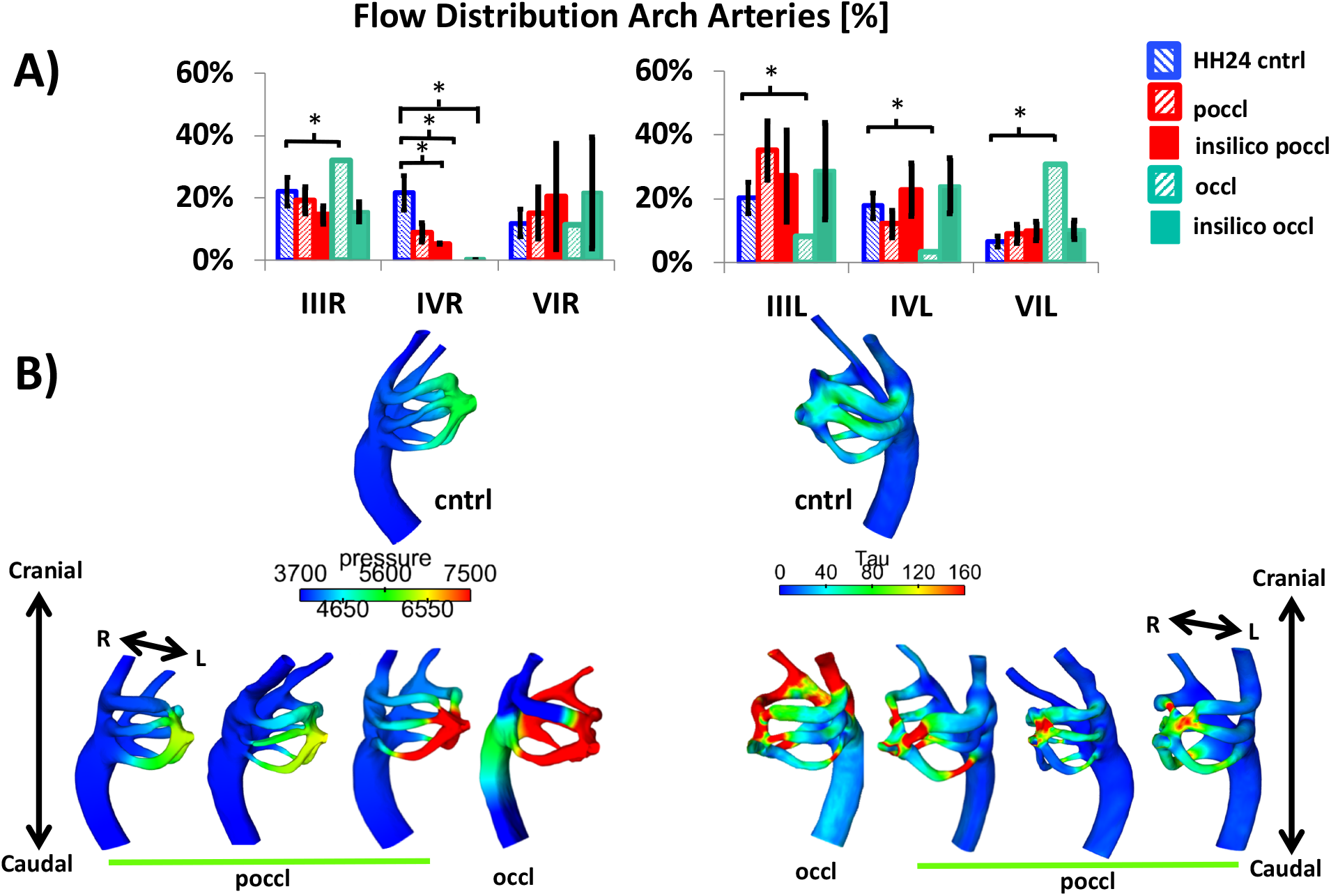
HH24 hemodynamic changes resulting from experimental vessel occlusion. (A) Flow distributions per arch (+/−s.e.) asterisks indicate a statistically significant change when a two tailed student t (poccl, in-silico occl/poccl) or one-sample t-test (occl) (B) Pressure and wall shear stress (dynes/cm^2^) at peak flow for a representative HH24 control embryo (top) and each of the partial occlusion geometries in which CFD was run. Although the geometries themselves have changed, partial occlusion pressure and WSS maps resemble that of HH24 control embryos (see [17] for range of variability). Only the full occlusion geometry’s pressure is so high that it cannot be fully dissipated along the arches and a high-pressure zone is carried over to the DoA. (N=5 cntrl, N=3 exp poccl, N =1 exp occl, N = 2 in-silico occl; insilico poccl).

Regardless of initial occlusion placement, structural analysis of HH24 occlusion geometries revealed a lengthening along the centerline of partial occlusion embryos, with a significant change observed in PAA III, IV and VI right, when compared to those of controls (Figure 3C). Arch arteries also became more circular in cross-section than their HH24 control counterparts, approaching the phenotype of an early HH26 embryo in both length and ellipticity, but maintaining a non-septated OFT (Figure 3A, B). HH24 arch artery area largely increases in PAA IIIL in order to compensate for HH18 IVR vessel occlusion (Figure 4A). Branch-splitting angle decreased for HH24 PAA IIIR and IIIL partial occlusion geometries when compared to that of HH24 controls (Figure 4B). When comparing HH24 partial and full occlusion branch splitting angles, the largest changes exist for the right DoA aorta branch-splitting angle and PAA IIIL.

**Figure 3.**
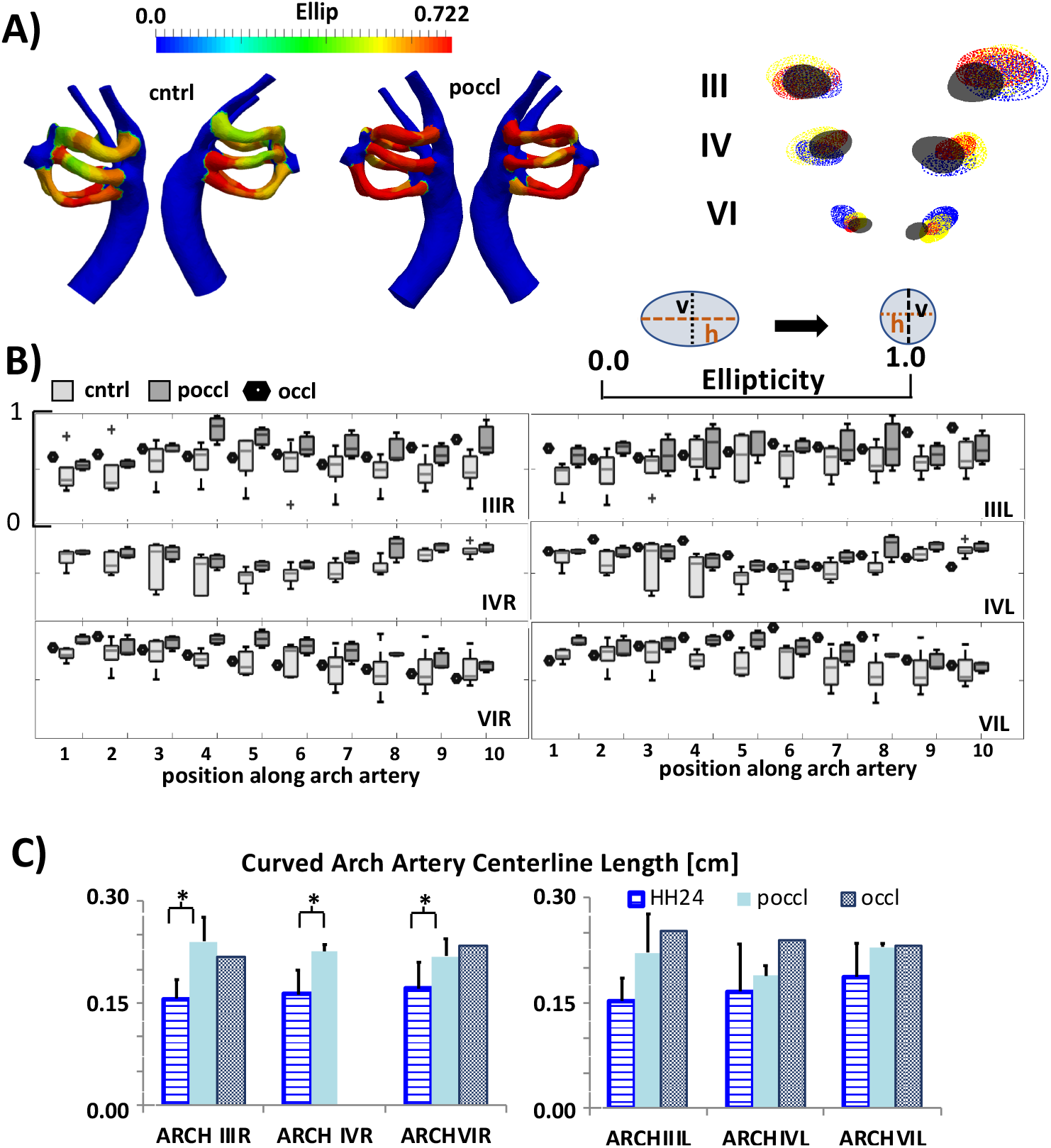
HH24 Occlusion Embryo Ellipticity and Length Changes Measured from 3D Reconstructions. Ellipticity changes for both the HH24 control and HH24 partial occlusion (poccl) geometries (A) Control and partial occlusion templates showing the average trend along with vessel cross-sections where control is displayed in black and overlayed over poccl cross-sections (B) Boxplots detail changes for each of the ten sections along the centerline of each arch (cntrl, poccl), single marker used for full occlusion (N =1). (C) PAA length per arch with asterisks indicating a statically significant difference when considering a two-tailed paired t-test. (N = 3, poccl, N = 5 cntrl). Occlusion geometries have more circular arch cross-sections and more elongated arches.

**Figure 4.**
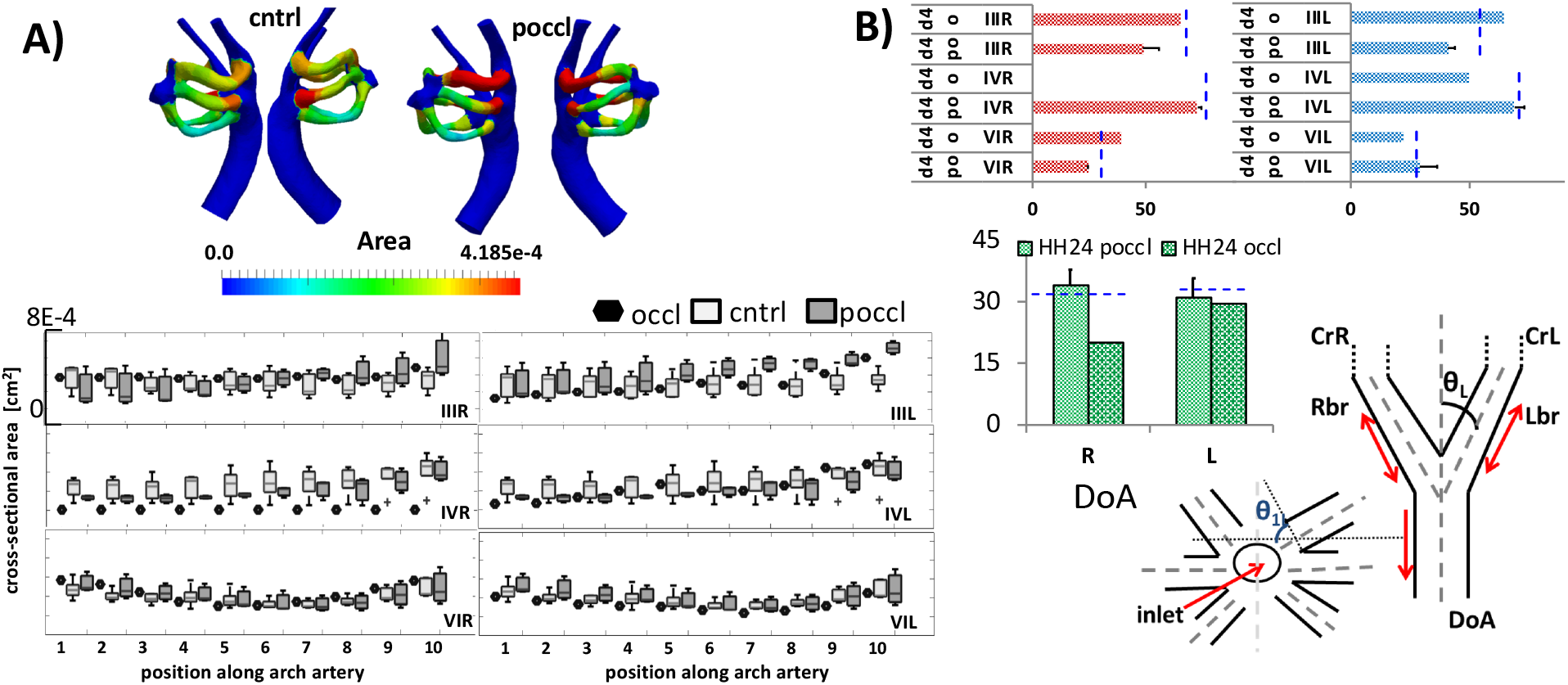
HH24 Occlusion Area and Angle Changes Measured from 3D reconstructions. (A)Cross-sectional area (CSA) for the control and poccl templates shown (top) with box-plots that detail the changes taking place along the centerline (‘T’ represents extreme bounds,’+’ outliers).Single marker used for full occlusion. The largest CSA changes are apparent in PAA III, particularly PAA IIIL and PAA IV. (B) branch-splitting angles with mean values for HH24 control displayed as dashed lines along corresponding arches for both HH24 partial occlusion (po) and full occlusion (o). The largest differences can be seen for PAA III, though no significant changes were found (N=5 cntrl, N=3 poccl, N =1 occl).

### Mechanical occlusion models of PAA development

To understand the observations described above, we developed an experimental-computational pipeline that begins by taking experimental-based computational models and re-distributing natural flow patterns through in-silico occlusion (Figure 5). In isolating the immediate flow and pressure redistributions, one can see how the embryo’s response differs from empirical predictions and begin to dissect the compensation mechanisms at work. Both full and partial occlusions were performed on HH18 and HH24 control geometries, which span the initial 24-hour remodeling period. Initial hemodynamic conditions were kept constant between control and in-silico occlusions, so as to highlight instantaneous flow redistribution when the vessel is not given time to adapt to changing hemodynamic forces. With only 5-7% of flow being re-distributed upon occlusion, changes in HH18 in-silico flow distribution were small (Supplemental Table 2A,B). Upon PAA IVR vessel occlusion, more flow is re-distributed to the left side of the embryo than the right in the full occlusion case (approximately 59% of flow re-direction is channeled to the left), and is distributed equally in the case of the partial occlusion. The corresponding changes in pressure and wall shear stress (WSS) as measured by numerical simulation are subtle (Figure 6B, top) with a slight increase in pressure magnitude visible at the aortic sac. Pressure and WSS magnitude in the left IV are slightly increased upon partial occlusion in both HH18 in-silico occlusion subsets. The results of both pressure and flow changes are summarized in Figure 6A in the form of resistance (See methods). HH18 PAA IVR resistance increases exponentially with vessel partial occlusion and is infinite in the case of full vessel occlusion when compared to that of controls, while resistance values for the other vessels remain largely unchanged. and full occlusion.

**Figure 5.**
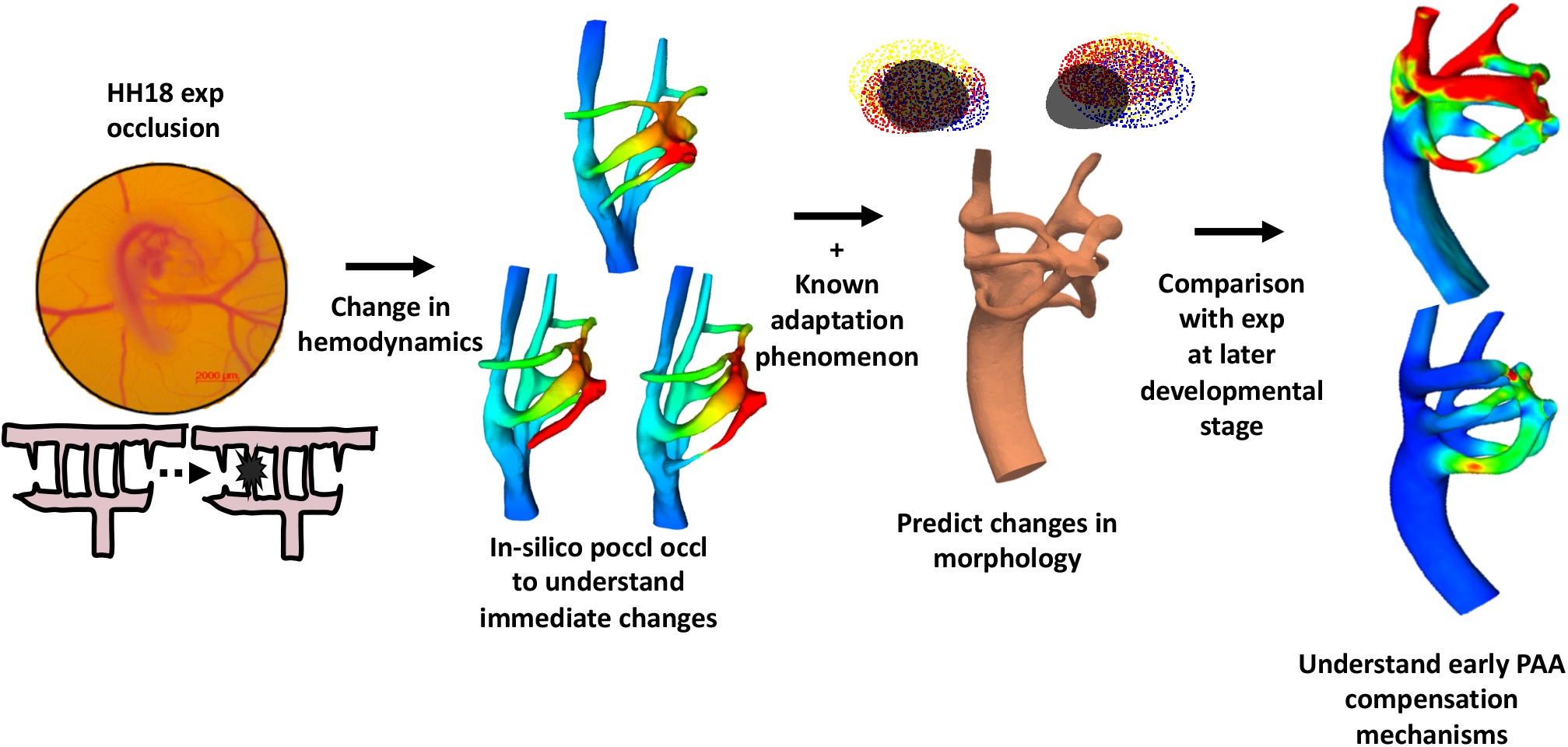
Combined experimental-computational pipeline for understanding hemodynamic-driven PAA flow adaptation. Highly targeted occlusion experiment led to changes in hemodynamics. In-silico occlusion experiments allowed for high resolution pressure and WSS maps. Combining immediate flow changes with known adaptation phenomenon allowed for the prediction of morphological changes. Theoretical predictions were compared with experimental geometries at later stages.

**Figure 6.**
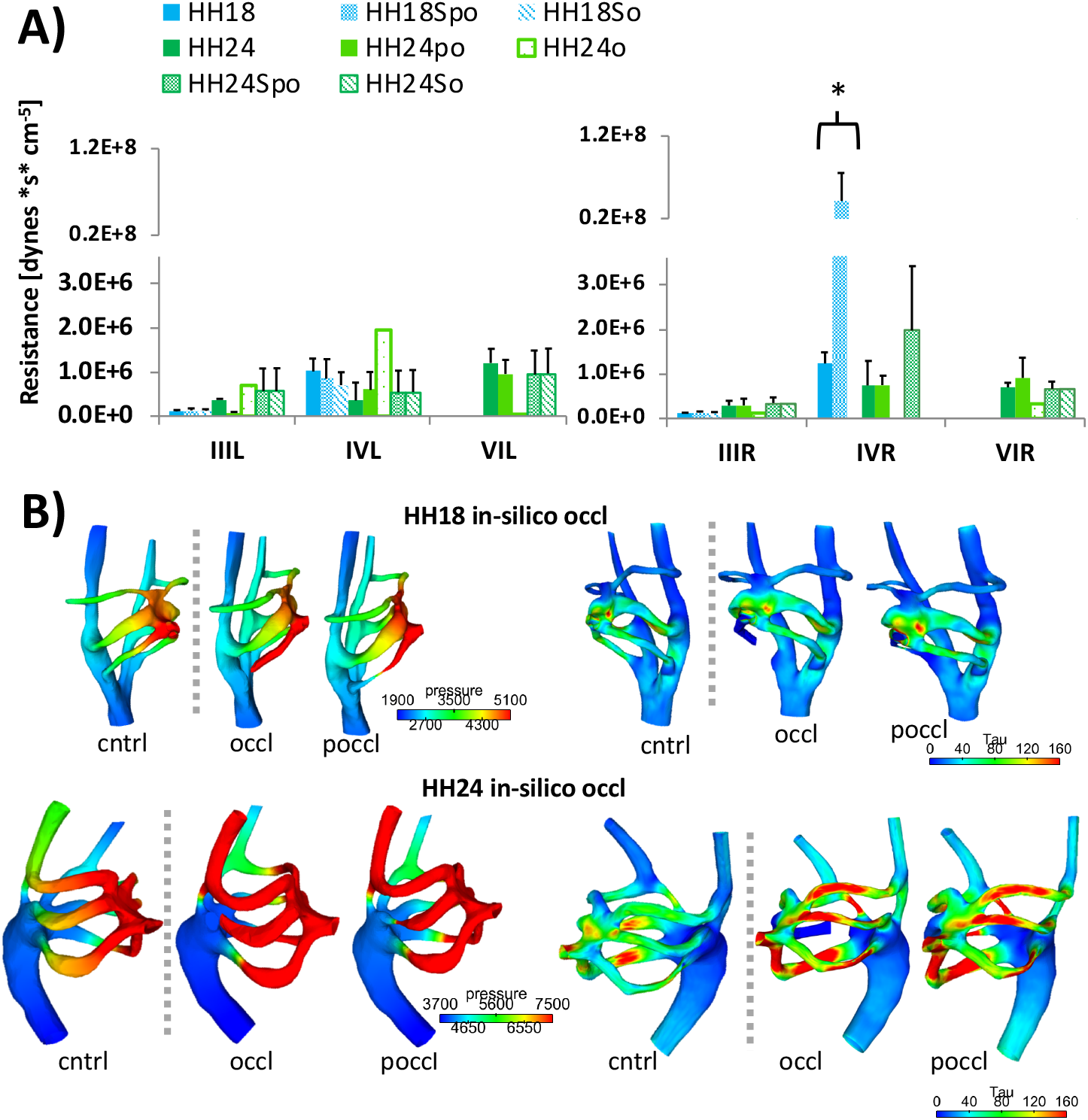
HH18 HH24 in-silico occlusion hemodynamic changes. (A) 0D resistance values for each arch per stage. The only significant difference found when comparing HH18 controls to in-silico occlusion or HH24 experimental occlusions to HH24 occlusions was for HH18 PAA IVR (HH18 to HH18So, two-tailed two-sample t-test). (B) Pressure and WSS (dynes/cm^2^) maps at peak inflow for a re-presentative HH18 and HH24 in-silico occlusion geometry. Note how HH18 in-silico occlusion effects are subtle due to the small amount of flow being re-distributed, while HH24 effects are more pronounced for both full a partial occlusion, through the difference between degree of occlusion is hard to distinguish. So, Spo = in-silico occlusion, partial occlusion, respectively. (N=5 cntrl, N=3 exp poccl, N =1 exp occl, N = 2 in-silico occl; insilico poccl).

Given the small value of flow re-distributed upon HH18 occlusion, two HH24 control geometries were occluded in-silico to assess the immediate effects of abnormal flow patterns on the more stable HH24 arch artery form in which no arches are in the process of growing in or regressing. Full and partial occlusions followed the same instantaneous flow re-distribution patterns, with PAA IIIL receiving 29% and 27% of the flow and PAA IVL receiving 24% and 23% of flow respectively in HH24 full and partial occlusion scenarios (Figure 2A). Changes in HH24 in-silico occlusion pressure and WSS maps (Figure 6B, bottom) are more pronounced than that of HH18 in-silico occlusions, but follow a similar pattern. Upon occlusion, pressure magnitude increased at the outflow tract inlet before dissipating cranially along the aortic sac, laterally along the IVR occluded or partially occluded arch as well as along the caudal VI arch arteries. Spikes in WSS increase in high pressure areas. These changes are summarized in the form of a dramatic increase in the resistance value of PAA IVR, small increase in PAA III (R &L) and PAA IVL. PAA VIR remain unchanged and PAA VIL slightly decreases. In both HH18 and HH24 in-silico occlusion scenarios, the embryo’s immediate response was to send additional flow to PAA IVL. 3D numerical simulations of experimental occlusion embryos reveled that in reality in-vivo flow distribution to PAA IVL decreased in both the experimental partial occlusion and full occlusion geometries, suggesting that other mechanisms may also be at play.

### Local hemodynamic changes dominate PAA growth & remodeling

Computational fluid dynamic simulations of HH24 experimental occlusion embryos (24 hours post occlusion) allowed us to investigate the evolution of hemodynamic flow patterns and associated forces. Experimental partial occlusion pressure and WSS maps resemble that of control embryos while the full occlusion falls within the same range but has a more elevated pressure and WSS magnitudes along the DoA (Figure 2B). The experimental occlusion embryo maintains the same general pressure dissipation pattern, but is unable to fully dissipate pressure before the arches converge at the DoA. To delineate the role of pressure and WSS as drivers of PAA remodeling, we quantified morphological changes from CFD-aided 3D renderings obtained from point-to point segment makers, as outlined in [17]. Statistically significant changes between relative position at HH18 to HH24 were found through linear regression of data obtained per arch. Linear regressions were performed using HH18 control data (N =5 embryos) or HH18 in-silico occlusion data (N =2 embryos) as time point 1 and either HH24 experimental occlusions (N = 3 partial occlusions, N =1 full occlusion) or HH24 in-silico occlusions (N = 2 in-silico occlusion) as time point 2 (Figure 7). HH18 to HH24 change in peak WSS magnitude leads to significant arch artery correlations for each of the arch artery pairs (Figure 7). For control embryos, the majority of WSS-morphology trends were associated with positive slopes and 30% were associated with a negative slope. Because flow through these arches are essentially parabolic (Poiseuille flow) and Poiseuille WSS magnitude is proportional to flow over radius cubed (Q/r^3^) or mean velocity over radius (ν_mean_/r), a positive slope suggests a change in flow/velocity is locally driving WSS change, while a negative slope implies the geometric feature is dominating local WSS change between stages. This indicates that following vessel occlusion, local changes in WSS were associated with remodeling along the arches, contrary to controls where decrease of WSS was rather related to flow.

**Figure 7.**
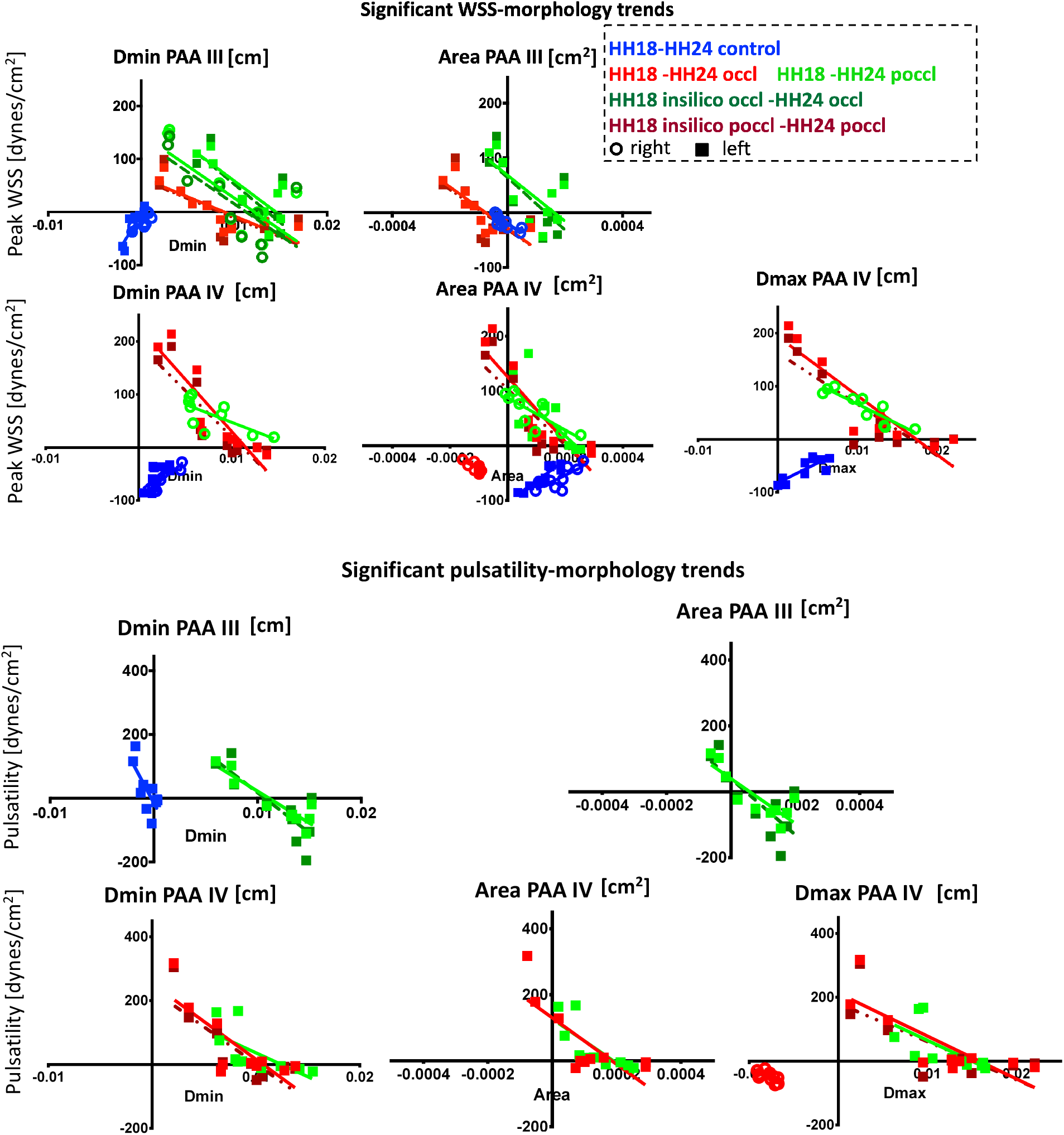
Occlusion driven geometric changes. Linear regressions plotted to highlight correlations between hemodynamic forces (peak WSS and pressure pulsatility) and area diameter changes that occur with occlusion insult. Axes show changes in x and changes in y values. Only significant trends are shown. (circles indicate right side, squares, left side). Control curves shown in blue, when significant. (N=5 cntrl, N=3 poccl, N =1 occl).

Pressure pulsatility (the difference between max and min pressure over a cardiac cycle) was also monitored for HH18 and HH24 embryos. Significant pulsatility-morphology trends were determined for PAA IIIL and IVL. Pulsatility-morphology trends were also associated with negative slopes. WSS-morphology trends for in-silico occlusion geometries were unsurprisingly dominated by local geometric changes rather than hemodynamics (Supplemental Figure 3), as the arch arteries were constrained to their control geometric configuration. Only hemodynamics was permitted to change upon in-silico occlusion; the vessels had no time or opportunity to remodel.

### Local occlusions affect pressure distribution to subsequent circulation

In addition to flow and pressure values within the arches themselves, vessel occlusion also affects flow and pressure distribution curves to subsequent circulation. Upon vessel occlusion, inlet pressure is affected (Figure 8), despite maintaining the same imposed flow across embryos, with the full occlusion model showing a higher spike than partial occlusion curves for their respective conditions (experimental, in-silico). Pressure curves entering the Windkessel bounds are largely the same for the left cranial branch, but are slightly elevated for in-silico occlusion and in-silico partial occlusion when compared to its control phenotype (Supplemental Figures 4, 5).

**Figure 8.**
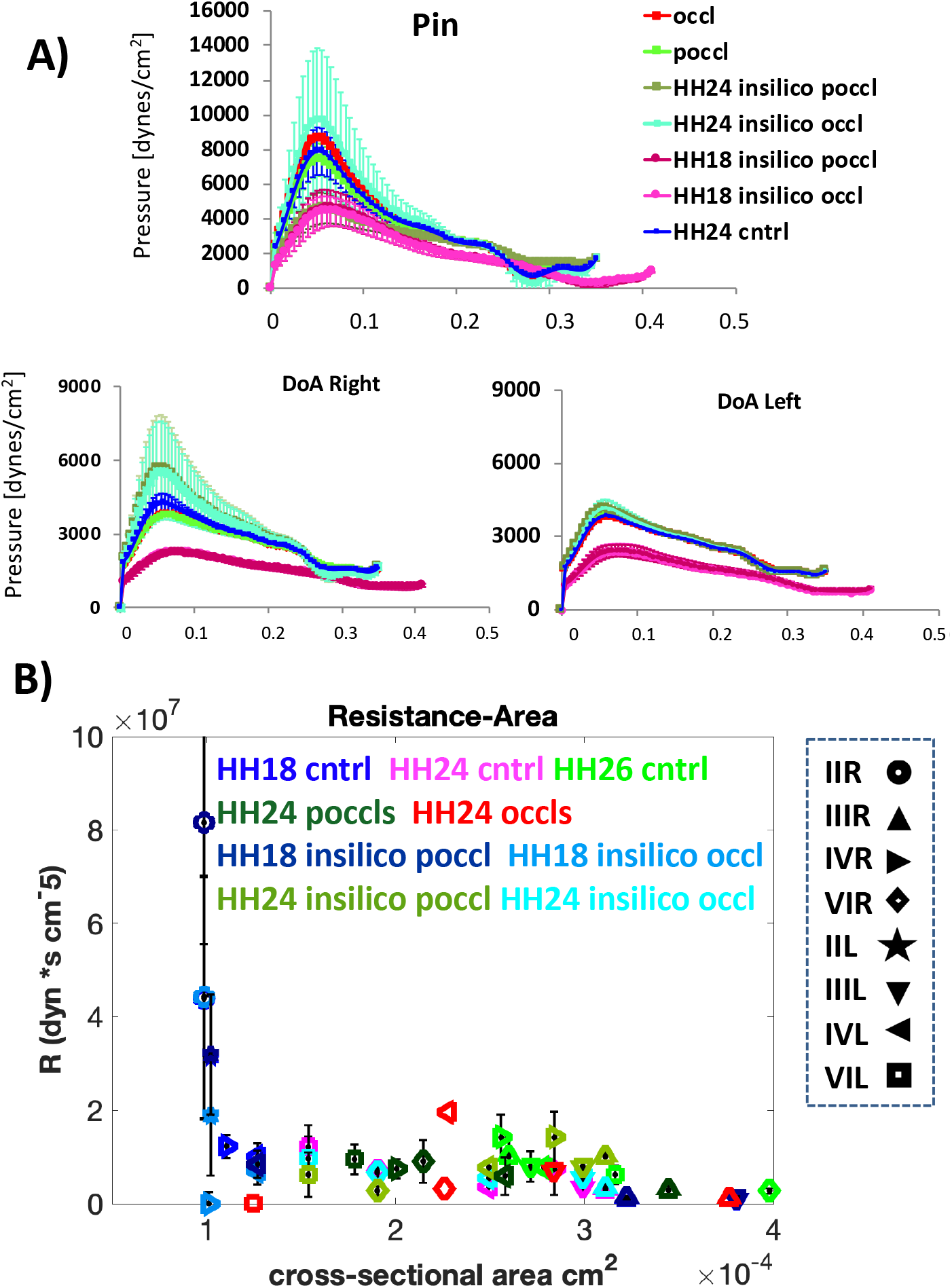
Pressure and functionality changes upon vessel occlusion. (A) 3D-0D resulting inlet pressure curves (top) and DoA branch (R & L) outlet curves for each of the occlusion conditions, both experimental and in-silico. Standard error bars are shown for each curve. (B) Resistance-area graph highlights vessel functionality. R-A axes show current x values with current y values. Standard error bars are shown in black. Note how HH24 in-silico occlusion predicted resistance values for three of the remaining five vessels for HH24 experimental partial occlusions. HH24 IIIL of the in-vivo full occlusion took on the same resistance and area values as HH24 IVR controls. PAA VIL of HH24 in-vivo full occlusion took on the same cross area as HH18IVL. (N=5 cntrl, N=3 exp poccl, N=1 exp occl, N=2 in-silico occl; insilico poccl).

90-10 cranial caudal flow distributions are largely maintained upon full and partial in-silico vessel occlusion of HH18 and HH24 embryos (Supplemental Table 3) despite bounds not being tuned to maintain the flow split. A slight deviation is seen the HH24 IVR in-silico occlusion, which has a 89-11 flow split. While changes in the caudal pressure curves are slight, changes in cranial pressures are much larger, particularly for HH18-1 full occlusion (Supplemental Figure 4, magenta). HH24 occluded and partially occluded embryos exhibit a much lower cranial pressure magnitude when compared to that of the control (Supplemental Figure 5). Very slight changes are seen in flow curves distributed to the rest of the embryo upon vessel occlusion.

### Instantaneous flow changes largely determine vessel functionality

The evolution of pressure and flow values along each arch artery determines its individual functionality, or each of flow throughout the cardiac cycle. Functionality changes are summarized in the form of resistance-area plots (Figure 8B). HH24 in-silico occlusion predicted resistance values for three of the remaining five vessels for HH24 experimental partial occlusions (PAAs VIL, IVL and IIR). Arch arteries underwent an increase or decrease in area to obtain these values. Of the arches not matched by instantaneous (in-silico) flow change predictions, PAA IIIL from the HH24 in-vivo full occlusion model obtained the same area and resistance values as PAA IVR from HH24 control embryos. PAA IIIL therefore assumed the profile of the normally flow dominant arch in control embryos. PAA VIR from HH24 in-vivo full occlusion model maintained the same CSA as HH24 in-vivo full occlusion PAA IVL.

A number of vessels maintained the same cross-sectional area (CSA) but different resistances values as peer vessels, emphasizing how CSA does not determine functionality. This is the case for PAA IVR in-silico occlusion and IIIL experimental occlusion. PAA VIL of HH24 in-vivo full occlusion assumed the same cross-sectional area as HH18IVL (control, in-silico occlusion/partial occlusion), indicating that IVL did not change morphology between HH18 and HH24. HH18 in-silico occlusion/partial occlusion PAA II (R & L) and PAA IVR share the same CSA value, but resistance values differ largely. These arches are regressing and growing in respectively.

### PAA adapts to abnormal hemodynamic signaling in two phases

Morphological changes identified in section one, indicate that the partial occlusion is causing the embryos to mature more quickly, elongating in size and displaying more rounded cross-sectional areas. Comparison of HH24 experimental occlusion hemodynamics and HH18/HH24 in-silico occlusion hemodynamics suggest a two – phase model for PAA’s response to altered hemodynamics (Figure 9) In the initial phase, immediately following vessel occlusion, a local increase in pressure propagates upstream to the aortic sac, leading to an increase in WSS. The aortic sac in turn increases in diameter, which causes a decrease in length of the PAAs which are pulled from their aortic sac inlets. The second phase, brought on by the stretching of the arches by the aortic sac, leads to an increase in axial stress, which is compounded by the flow induced asymmetry, culminating in the overall increase in longitudinal length experienced by the arches. Concurrent with the phase 1’s local increase in pressure, is a decrease in WSS experienced by the occlusion branch which leads to a shrinking or altogether elimination of the arch and adds to the flow induced asymmetry. In this way, natural growth mechanisms are overridden and the would be presumptive aortic arch is prevented from becoming the dominant arch of the aorta.

**Figure 9.**
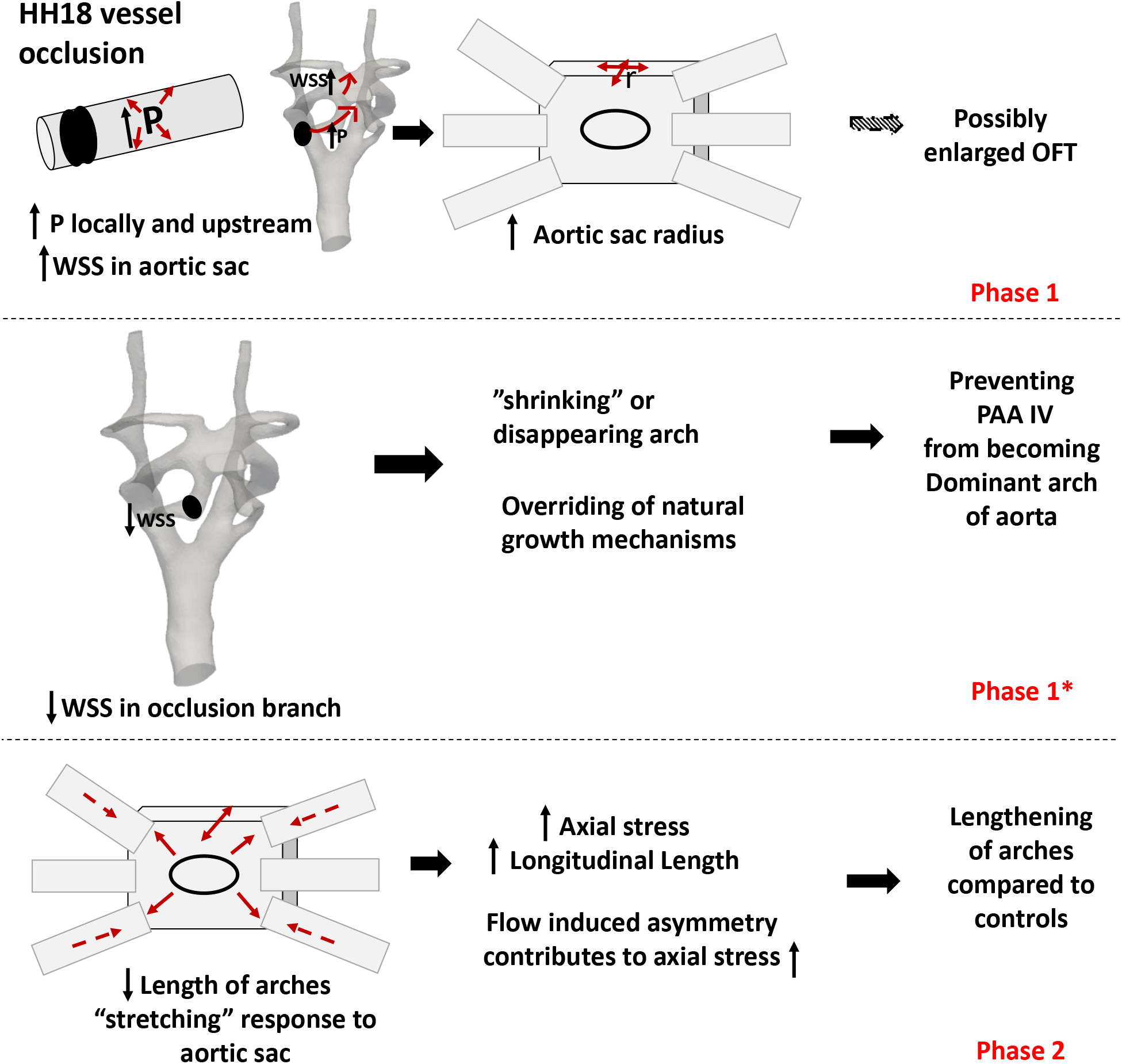
Two-phase vessel adaption mechanism following vessel occlusion. Local and upstream pressure changes lead to WSS changes that ultimately shape the structural changes seen in experimental occlusion embryos. Changes in aortic sac diameter lead to increased axial stress and accounts for the longitudinal lengthening of the arch arteries.

### Small initial insult evolves into larger structural deformation

In order to underscore the importance of the remodeling presented throughout this paper in embryos 24-hours post PAA IVR occlusion, a partial occlusion embryo, Exp-poccl-10, was allowed to develop to HH29 (day 6), when the aortic and pulmonary trunks are clearly distinguishable, the left IV arch artery has regressed, and the aortic trunk has begun to cross under the pulmonary trunk. At HH24 the partial occlusion embryo’s stroke volume, cardiac output and ventricular ejection fraction differ slightly from that of HH24 control embryos; only HH24 partial occlusion OFT diameter was statistically significant (two-tailed t-test). By HH26 both the OFT diameter and ventricular ejection fraction of Exp-poccl-10 were statistically different from that of HH26 controls when using a one-sample T-test. By HH29 morphological differences are readily visible with the poccl geometry displaying hyper-enlarged arches and abnormal rotation of the OFT (Figure 10, Supplemental Figure 6).

**Figure 10.**
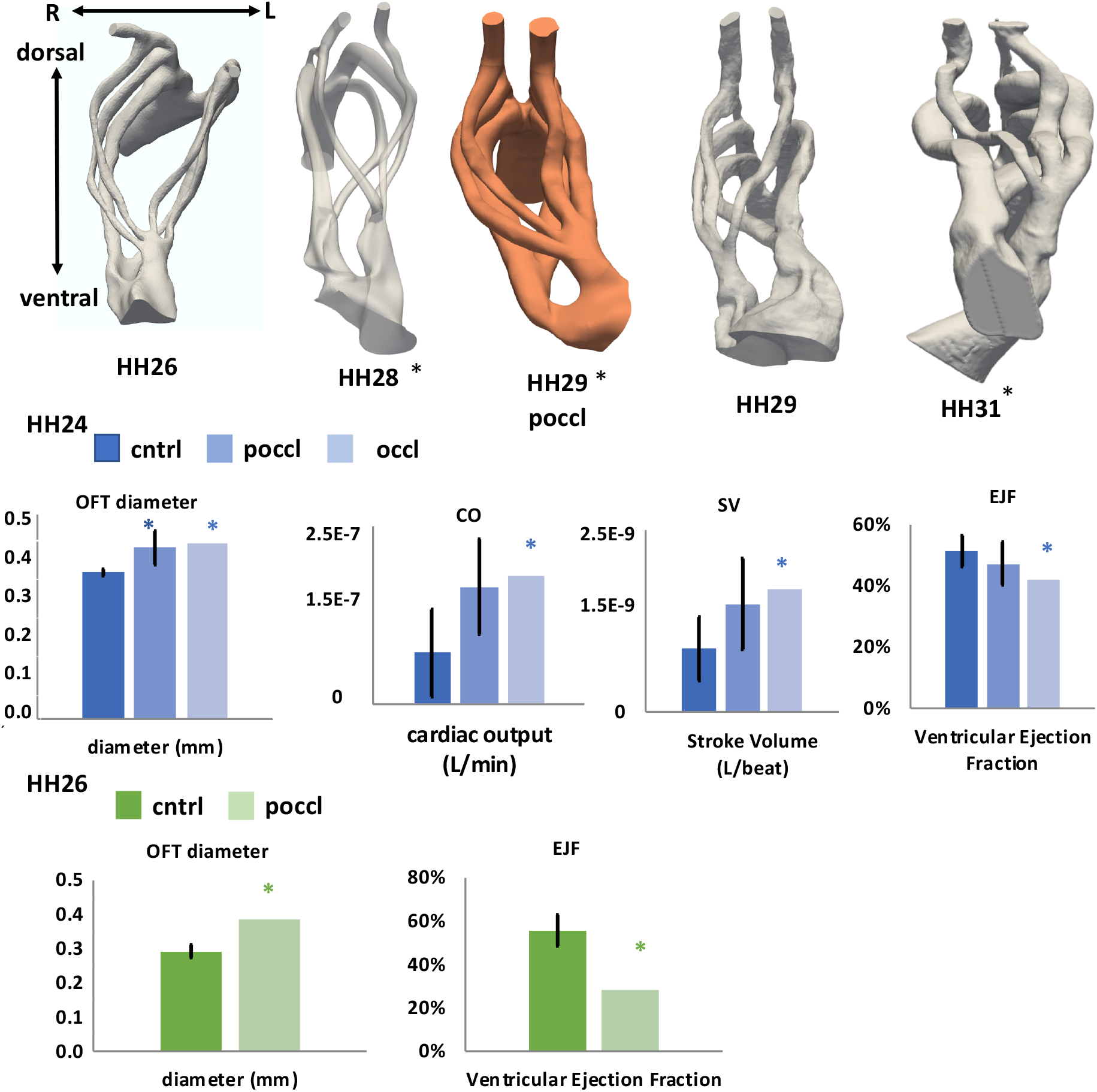
Later stage partial occlusion phenotype. (A) Top down view of PAA manifold of HH26 -HH31 embryos with controls in grey and partial occlusion in peach. Asterisks indicate that the embryo is left-sided. Geometries are not shown to scale. Of note is the presence of six arch arteries in the HH29 partial occlusion manifold, when five are present in control counterparts. The enlarged arch arteries of the aortic and pulmonary trunks run parallel to each other along the dorsal-ventral axis in contrast to their control counterparts whose aorta and pulmonary trunks are offset allowing for the aortic trunk to pass under the pulmonary trunk. B) HH24 diameter cardiac output, stroke volume, ventricular ejection fraction (n = 5 control, n = 3 poccl, n = 1 occlusion). Statistically significant differences were found for OFT diameter changes and when comparing the occlusion phenotype to controls using a one sample t-test. The highlighted poocl from (A) is among poocl values. (C) HH26 diameter and ejection fraction for HH26 embryos (n = 5 control, n = 1 poccl). Statistically significant differences were only found when comparing the occlusion phenotype to controls using a one sample t-test. The highlighted poocl from (A) is among poocl values.

## Discussion

From HH18 – HH24, the arch arteries undergo many changes. These changes are the result of inlet flow stream distributions, 3D aortic sac and aortic arch geometries, and local vascular biologic responses to spatial variations in WSS [1]. Our major findings concern how mechanical interaction between fluid flow in the arches and structural configuration of the vessels influence cardiac outflow morphogenesis. In particular, we found that a local increase in pressure accompanied by a decrease in WSS along the occluded vessel ultimately resulted in observed vessel elongation and CSA changes. The early embryo adapted its vasculature using known vascular adaptation mechanisms normally associated with the mature vessel. In the mature vascular system, the vascular network works to maintain constant homeostatic stress which consists of shear, axial and circumferential stress along the vessels [18]–[20]. The WSS stimulus leads to diameter changes in response to flow rate [21]. In a 1980 canine carotid artery study, vessel diameter was shown to increase with increased flow load and vice-versa (Kamiya and Togawa) as a way of bring shear stress down to control levels. Similarly, a 1987 study observed vessel dilation and no hypertrophy in response to increased flow in the iliac artery of monkeys [22]. HH18 experimental occlusion embryos were also able to adapt their diameters, over the span of 24 hours, to maintain a desired stress state.

The HH18 time point is a particularly critical stage of cardiovascular development as it precedes migration of neural crest cells down the pharyngeal arches. Following formation of arch arteries III, IV, VI, neural crest cells seed the arch arteries, covering their endothelial sheath and making up their ectomesenchyme [23]. A subset of these cells reach the outflow tract cushions by HH23. Cardiac neural crest cells (CNCCS), along with a shelf of tissue from the aortic sac, form the aortopulmonary septation complex and initiate septation [24]. Disruption of this process due to altered mechanical loads within the vessels would lead to a variety of cardiac abnormalities. CNCC ablations are associated with failure of arch arteries three, four (right), and six to develop to the proper size [25]. There is a loss of bilateral symmetry, with uncommonly small or collapsed arch arteries on one side and unusually large arteries on the opposite side [26]. Many of these phenotypes can be seen in the experimental occlusion geometries found in Supplemental Figure 1. Typically, asymmetrical remodeling begins at HH28/E12.5[27], [28], with the left fourth arch artery regressing in the chick embryo and the right fourth in the mouse embryo. The experimental late stage (HH29) partial occlusion embryo, Exp-poccl-10, did not undergo this asymmetric remodeling. Its phenotype approaches that of the *Pitx2δc* mutant hearts with transposition of the great arteries, where the aortic and pulmonary trunk remain parallel. *Pixt2* regulates left-right asymmetry to asymmetrically developing organs and is associated with patterning of the OFT myocardium[29]–[31]. This myocardium is derived from the pharyngeal mesoderm. Between Stages HH21 and HH30, Carnegie stages 15 and 19, the OFT-PAA junction undergoes rotation in the counterclockwise direction[32], [33]. Bajolle et al showed that abnormal rotation of the myocardium wall of the OFT in early embryos (E11.5/HH24) was associated with abnormal positioning of the great vessels (2006). Here we show that this defect may be the result of abnormal flow patterning originating from the PAAs themselves.

This study demonstrates that abnormal PAA hemodynamics can precipitate abnormal cardiac function given the correct timing and location of injury. In their 1997 study, Kirby et al altered arch artery patterning through molecular modification of premigratory neural crest cells. Despite affecting arch artery patterning during the same critical window of development, HH18-HH24, Kirby et al showed that altered patterning of the aortic arches does not necessarily result in abnormal cardiac development. Kirby et al were able to cause persistence of a transient arch, abnormal regression of a mature arch, and the appearance of an entirely new arch. In all cases the IVR PAA was still permitted to function as the dominant arch artery and the PAA defect was resolved by a respecification or reprogramming of existing arches. In the case of the disappearing PAA IIIR, PAA IVL supplied the necessary pathways and the normally transient PAA V remained patent through the transition [34]. These results underscore the fact that location as well as the timing of the initial insult are critically important to malformation risk. In their 2007 study, Yashiro et al. demonstrated that differential flow patterns and abnormal PAA remodeling can result from abnormal spiraling of the OFT. Following Pixt2 induction of abnormal OFT morphology, differential flow patterns led to abnormal molecular expression patterns and abnormal PAA VI patterning in E11 −11.5 mice (HH18-HH24 equivalent) [35]. Here, we show the connection between abnormal PAA IV patterning and OFT morphology, with the defect originating from within the PAAs themselves.

This study demonstrates a role for WSS in the initiation and propagation of cardiac defects of the OFT. Following arch artery occlusion, local changes in WSS are associated with geometric remodeling along the entire PAA manifold. Wall shear stress drives defect propagation which can result in enlarged OFT diameters, irregular regression patterns and abnormal rotation of the OFT. Though our group has previously studied the effects of arch artery occlusion on HH18 and HH24 embryos [3], the experimental results were not considered in 3D, which allows for a more thorough investigation of pathologies [36]. In our 2018 study we detailed morphological and hemodynamic changes in control HH18 HH24 and HH26 embryos [17]. In light of this work we can quantify the deviation from normal exhibited in HH24 embryos, upon HH18 vessel occlusion. Using our experimental occlusion pipeline (Figure 3), we are able to detail the phases of vessel growth and adaptation taking place. By examining embryo-specific geometries in 3D, placing them back in the context of the body (through physiologically relevant bounds), and comparing them to that of in-silico occlusion embryos, we distinguish immediate changes from that of longer-term adaptation to changing hemodynamic conditions and show how the former influences the later. This work can be used to identify new genes and biochemical compounds that affect cardiac outflow morphogenesis.

## Methods

### Creation of In-vivo Occlusion Models

Embryos were experimentally occluded as previously described [3]. Briefly, HH18 embryos were open cultured. Embryos with smaller IVR arch arteries were specifically chosen to facilitate faster occlusion experiments. Selected embryos were injected with Texas red dextran (70,000 MW, neutral Sigma-Aldrich D1830) diluted in Earle balanced salt solution (5% w/v) so that their vasculature could be visualized by way of two-photon microscopy. A custom built two-photon excited fluorescence microscope with a separate femtosecond pulsed photoablation laser was used to perform targeted vessel disruption. The photoablation laser consisted of 1-kHz high-pulse-energy Ti:Sapphire amplified laser system with 50-fs pulse duration (Legend-USP, Coherent, Santa Clara, CA, 800-nm central wavelength). Photodisruption was controlled and confined to the volume focused by two-photon microscopy optics.

### Ultrasound processing and generation of computational inlet flow curves

Outflow tract (OFT) velocity and that of the three paired pharyngeal arch arteries were measured using B-mode guided Doppler Ultrasound (Vevo770 and Vevo 2100, Visualsonics, Inc.) and converted into flow rate, as explained in [17]. A Poiseuille OFT profile was assumed and maximum velocity converted to flow. The mean calculated flow was fed to each embryo via a steady-state simulation and the ratio between flow rate and velocity calculated in each of the individual arches. A new inlet flow, Qinlet, was then calculated to be the sum of flow in each of the individual arches as previously described [17].

### In-silico geometry preparation and 3D-0D flow modeling

Embryo-specific 3D geometries of HH24 PAAs were generated from experimental occlusion geometries by importing nano-CT images into MIMICS (Materialise, Louvain, Belgium) and 3MATICs (Materialise, Louvain, Belgium), as outlined in [17]. Geomagics Studio 10 (Geomagic Inc., Durham, NC) was also used for the preparation of 3D geometries for CFD. All embryos were scaled by a factor of three to account for the difference between dehydrated and native vessel size. Blood was treated as an incompressible Newtonian fluid with constant hemodynamic properties (ρ = 1060 kg/m^3^, μ = 3.71 × 10^−3^ Pa.s) and rigid, impermeable vessel walls were assumed with no slip boundary conditions. Flow was simulated by solving the Navier-Stokes equations, on a high-resolution unstructured grid with finite-element numerical treatment (solved using the finite element library FELiSce) (gforge.inria.fr/projects/felisce) [37]. Grid sensitivity analysis was conducted on a control PAA model for each day in order to ensure consistency and reliability of the numerical solutions for all simulations presented in this study, beyond which resulting mass-flow redistributions were insensitive to further Cartesian grid refinements.

#### boundary conditions

A natural spatial velocity profile was imposed at the inlet. To obtain this an auxiliary steady Stokes equation, with natural boundary conditions at the outlets, was solved first. The resulting inlet velocity profile was subsequently scaled at each time-point to match the measured flow-rate Qinlet. RCR Windkessel models were imposed at the outlets to assure the distribution of the cardiac output to dorsal aorta (DoA) and cranial vessels maintains a 90-10 flow split over the course of one cycle as summarized in the 0D section.

The differential equation representing the RCR circuit is

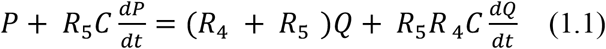

Where R_4_ is the proximal resistance, R_5_ is the distal resistance, C represents capacitance, P is pressure and Q is flow.

### Creation of In-Silico Occlusion Models

In silico occlusion models were made from two HH18 and two HH24 pharyngeal arch artery systems. The two HH18 control references (HH18-1 and HH18-5) were chosen to represent the wide range of variability seen across HH18 embryos [17]. HH18-1 has a rapidly regressing right and left lateral second arches as well as a developing right and left lateral forth arch, while HH18-5 has the largest right and left lateral second arch, as well as the largest right and left lateral fourth arch, seen in our HH18 control subset (Supplemental Figure 7B). Similarly for HH24, two embryos (HH24-1 and HH24-4) were chosen because of pressure range and because HH24-1 exhibited right flow dominance, while HH24-4 exhibited left flow dominance[17]. A full occlusion and a partial occlusion were made for each of the four reference geometries, bringing the number of in-silico occlusion geometries to eight. Partial occlusions were made to mimic those seen in experimentally occluded embryos, either through two-photon microscopy (HH18 embryos) or their subsequent nano-CT reconstructions from images taken 24 hours post vessel occlusion.

### Statistical Analysis

Morphological and hemodynamic changes were compared qualitatively and quantified when possible. Results were summarized in the form of mean and standard deviation values over the course of one cardiac cycle. Two-tailed two-sample t-tests were used where appropriate with p < 0.05 denoting significance. A one sample t-test was also used when comparing a single occlusion sample to that of controls. Linear regressions were performed. Statistical comparisons were made through the use of GraphPad Prism (GraphPad Software, Inc San Diego, CA) statistical software. Linear regressions were performed on hemodynamic versus geometrical parameter graphs.

## Abbreviations

HH: Hamburger Hamilton
PAAs: pharyngeal arch arteries
OFT: outflow tract
DoA: Dorsal aorta
poccl: partial occlusion
occl: occlusion
WSS: wall shear stress

## Acknowledgements

We thank the Cornell BRC imaging facility and Randolph Linderman (reconstructions). This work was funded by NSF GRFP, NSF GROW, an Alfred P. Sloan Foundation fellowship, CMMI-1635712, an Inria International Internship grant and by the National Institutes of Health (HL110328, S10OD012287 and S10OD016191).

